# The “DDVF” motif used by viral and bacterial proteins to hijack RSK kinases evolved as a mimic of a short linear motif (SLiM) found in proteins related to the RAS-ERK MAP kinase pathway

**DOI:** 10.1101/2024.08.08.607128

**Authors:** Martin Veinstein, Vincent Stroobant, Thomas Michiels, Frédéric Sorgeloos

## Abstract

Proteins of pathogens such as cardioviruses, kaposi sarcoma-associated herpes virus, varicella zoster virus and bacteria of the genus *Yersinia* were previously shown to use a common “DDVF” (D/E-D/E-V-F) short linear motif (SLiM) to hijack cellular kinases of the RSK (p90 ribosomal S6 kinases) family. Remarkable conservation of the SLiM docking site in RSKs suggested a physiological role for this site. Using SLiM prediction tools and AlphaFold docking, we screened the human proteome for proteins that would interact with RSKs through a DDVF-like SLiM. Using co-immunoprecipitation experiments, we show that two candidates previously known as RSK partners, FGFR1 and SPRED2, as well as two candidates identified as novel RSK partners, GAB3 and CNKSR2 do interact with RSKs through a similar interface as the one used by pathogens, as was recently documented for SPRED2. Moreover, we show that FGFR1 employs a DSVF motif to bind RSKs and that phosphorylation of the serine in this motif increases RSK binding. FGFR1, SPRED2, GAB3 and CNKSR2 as well as other candidate RSK binders act upstream of RSK in the RAS-ERK MAP kinase pathway. Analysis of ERK activation in cells expressing a mutated form of RSK lacking the DDVF-docking site suggests that RSK might interact with the DDVF-like SLiM of several partners to provide a negative feed-back to the ERK MAPK pathway. Thus, through SLiM mimicry, pathogens not only retarget RSKs toward unconventional substrates but also likely compete with human proteins to alter the regulation of the RAS-ERK MAP kinase pathway.

**Author Summary:** Short linear motif (SLiM) are 3 to 10 amino acid-long protein sequences that can mediate the interaction with other proteins. We previously observed that highly unrelated pathogens, including viruses and bacteria, convergently evolved to hijack cellular enzymes of their host, through a common SLiM. In this work, we tested the hypothesis that the SLiM found in proteins of pathogens evolved to mimic a SLiM found in human proteins that regulate the cellular enzymes through the same interface. Protein-protein interactions mediated by SLiMs are often, low-affinity, transient interactions that are difficult to detect by conventional biochemical methods but that can nowadays be predicted with increasing confidence by artificial intelligence-based methods such as AlphaFold. Using such predictions, we identified several candidate human proteins and we confirmed experimentally that these proteins interact with the cellular enzymes the same way as pathogens’ proteins do. Identified proteins belong to the well-known RAS-ERK MAPK pathway which regulates important functions of the cell, suggesting that pathogens evolved to hijack this MAPK pathway by SLiM mimicry. By doing so, they can both dysregulate cellular physiology and hijack cellular enzymes to their own benefit.

## Introduction

As obligate cell parasites, viruses have evolved a number of ways to exploit or to interfere with key pathways in the cell, including the mitogen-activated protein (MAP) kinase pathways, in order to promote their own replication or to evade immune responses (1, 2). RSKs (p90 Ribosomal S6 Kinases) form a family of four closely related serine/threonine kinases which act downstream of the RAS-ERK (Rat sarcoma virus / Extracellular signal-Regulated Kinase) MAP kinase pathway and regulate essential cellular processes such as growth, proliferation, survival, motility, and immunity (3). Since these kinases play pivotal roles in cell physiology, tight regulation of their activity is essential (4–7) and dysregulation of this signaling pathway, including that of RSK kinases, has been associated with developmental disorders and cancer (RAS-ERK-MAPK: (8–10), RSKs: (11–15)). To ensure tight regulation, serine/threonine kinases commonly employ surface-located amino acids to interact with partners. These partners can be preferential substrates of the kinase or regulatory proteins that modulate kinase activity through catalysis, relocalization, or allostery (16). The recruitment of binding partners may occur via short linear motifs (SLiMs) usually present in disordered regions of protein sequences. SLiMs play a significant role in the regulation of the human interactome (17), and the disordered regions from where they arise are mutated in more than 20% of human pathologies (18). However, their low affinity and amino acid sequence degeneracy make them challenging to identify. To predict SLiM binding in docking grooves, Verburgt *et al.* and others have shown that AlphaFold-Multimer (19) exhibits remarkable performance, allowing to predict potential interactors at docking sites (20, 21).

SLiMs, which easily arise in viral protein sequences due to their rapid evolution, can be exploited by viruses as subversion strategies (22, 23). We previously reported that unrelated pathogens, such as RNA and DNA viruses (cardioviruses and herpesviruses) as well as bacteria (from the genus *Yersinia*), convergently evolved to use of a short linear D/E-D/E-V-F motif (further referred to as “DDVF”) to hijack host protein kinases of the RSK family (13, 14, 24). The DDVF motif encoded by these pathogens interacts with a region encompassing the KAKLGM residues, situated in a surface-exposed loop of the RSK kinases. Surprisingly, although surface-exposed loops and pathogen-targeted regions of proteins often undergo rapid evolution, the KAKLGM residues of the SLiM-docking site remained remarkably conserved among all four isoforms of human RSK as well as across evolution. This suggests that microbial proteins evolved to target an essential RSK regulation site and raises the possibility that some cellular proteins might interact with RSK through the same interface (13, 14). This work thus aimed at identifying such cellular RSK partners to test whether pathogens acquired this DDVF sequence by SLiM mimicry.

## Results

### 1. Screening for candidate human proteins that bind RSKs through a DDVF motif

Previous findings showed that some proteins from pathogens evolved a DDVF motif that binds and regulates RSKs, effectively subverting a well-conserved region of the kinase. This prompted us to explore whether cellular proteins might interact with RSKs using the same interface (13). To this end, we screened the human proteome for the presence of D/E-D/E-V-F motif using SlimSearch4 (http://slim.icr.ac.uk/slimsearch/) (25) and found 222 hits including several known RSK partners (**Table S1A)**. These protein candidates were further probed for RSK binding through AlphaFold-multimer machine learning structure prediction (19). To this end, peptides encompassing the DDVF motif were in silico-folded with a fragment from the human RSK2 N-terminal kinase domain (21). Out of 222 predictions, 65 were predicted, by at least one AlphaFold model to bind RSK2 in the KAKLGM docking groove in the same configuration as the ORF45 protein encoded by KSHV (14) (**Table S1C)**. Of these, twelve proteins were described as known RSK interactors according to Biogrid and IntAct interaction databases, albeit with unknown binding interfaces (26, 27). Remarkably, 10 out of 12 (∼85%) known RSK partners docked in RSK according to AlphaFold prediction against 65 out of 210 (∼31%) for unknown partners further corroborating the interest of such AlphaFold screens.

### 2. SPRED2 and GAB3 DDVF motifs mediate RSK interaction

Among the 65 predicted RSK-interacting proteins identified above, we identified SPRED2 and GAB3 as candidates carrying the DDVF motif in a putative disordered region predicted to dock on RSK by AlphaFold-multimer in 5 out of 5 models (**Supporting information S1C**). Despite their location in an unstructured region typically prone to variation, the DDVF motifs of SPRED2 and GAB3 are remarkably conserved throughout evolution supporting the assumption that they are functionally important (**Figure 1A-B**). Despite SPRED2 showing a high degree of sequence similarity with its isoforms, the DDVF motif present in SPRED2 is not found in SPRED1 or SPRED3. Similarly, the DDVF motif in GAB3 is absent in GAB1, GAB2, and GAB4. (**Supporting Figure S1A-B**).

**Figure 1:**
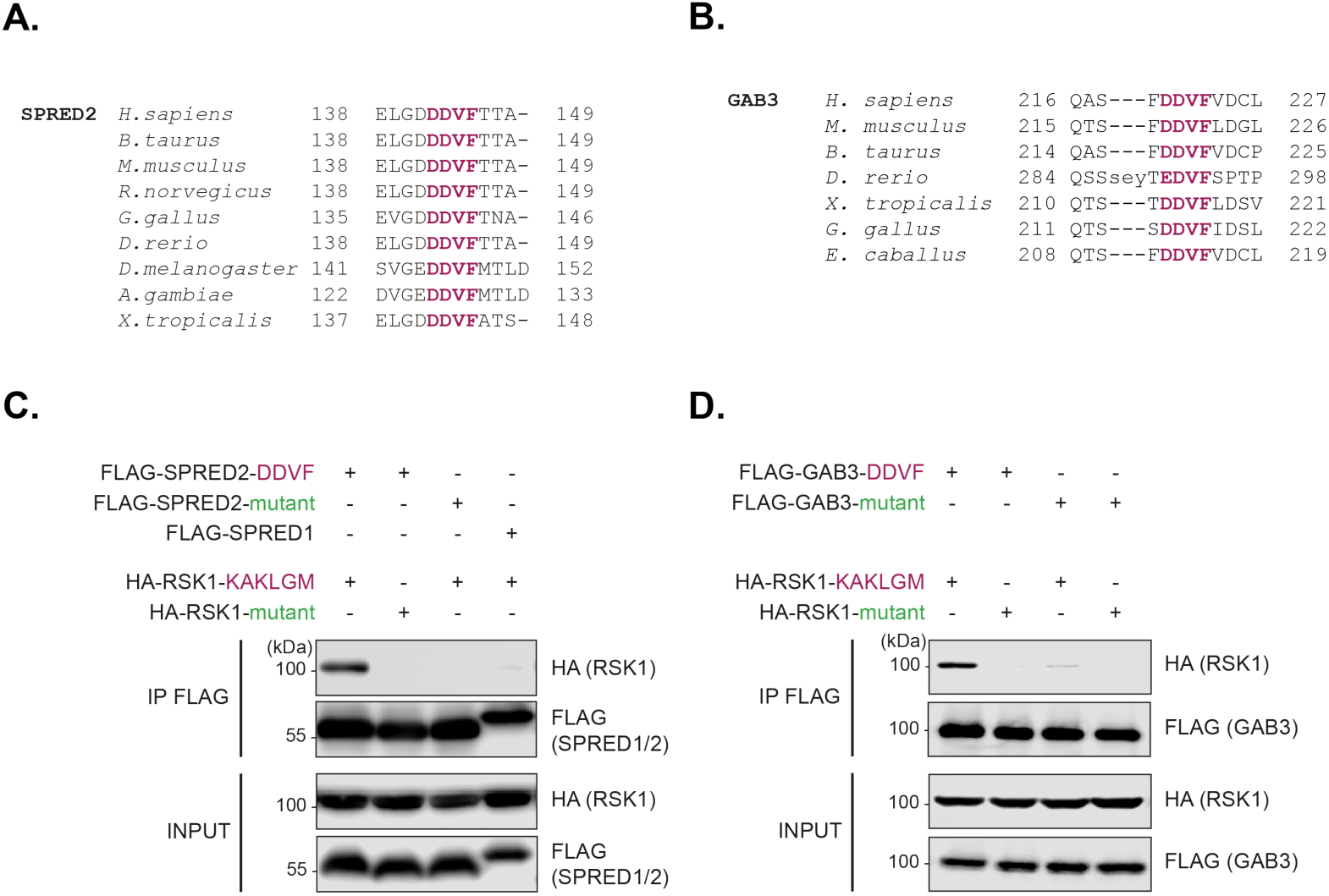
Highly conserved DDVF motifs in SPRED2 and GAB3 mediate the interaction with RSK1**. A-B.** Conservation of SPRED2 (A) and GAB3 (B) DDVF motifs (purple) across evolution **C-D.** Immunoblots showing the detection of wild type (KAKLGM) or mutated (KSEPPY) HA-RSK1 and FLAG (SPRED or GAB3 variants) in lysates (INPUT) of transfected HEK293T cells or after co-immunoprecipitation (IP FLAG) with various FLAG-SPRED variants (SPRED2-DDVF, SPRED2-DDV<u>A</u> mutant, or SPRED1), (n = 3) (C) or FLAG-GAB3 variants (GAB3-DDVF or GAB3-DDV<u>A</u> mutant), (n = 2) (D).

We used co-immunoprecipitation experiments to probe for an interaction of SPRED2 and GAB3 with RSK1. Controls included DDVF-to-DDVA mutations in the SLiM of SPRED2 and GAB3, as well as a KAKLGM-to-KSEPPY mutation in the SLiM docking site of RSK1. Both types of mutations were previously shown to dramatically reduce the interaction between microbial proteins and RSKs (13). As shown in **Figure 1C**, HA-RSK1 unambiguously co-immunoprecipitated with WT but not mutant FLAG-SPRED2 and FLAG-GAB3.Accordingly, HA-RSK1 did not co-immunoprecipitated with FLAG-SPRED1, which lacks a DDVF motif. Moreover, WT but not mutant RSK1 co-immunoprecipitated with FLAG-SPRED2 and FLAG-GAB3, suggesting that these proteins interact with RSK through the same DDVF:KAKLGM interface as the one used by microbial proteins. While this work was in progress, an elegant work by Lopez *et al.* (6) also showed the interaction of SPRED2 and RSK and provided a solution structure of the DDVF:RSK interface. This structure largely confirmed the similarity of the interface formed between RSK and SPRED2 and that formed between RSK and pathogens’ proteins, with the phenylalanine of the DDVF motif inserting in a hydrophobic pocket of RSK, at the level of the KAKLGM sequence (6).

### 3. Interaction of FGFR1 with RSK involves a DSVF motif which is subjected to regulation by phosphorylation

Our data (13), those of Alexa *et al.* (14) and Lopez *et al.* (6) suggested a critical role of the VF residues within the DDVF motif for RSK binding but also of nearby acidic residues thought to increase affinity through electrostatic interactions. The presence of two aspartic residues surrounding the VF motif were not absolutely essential, one being sufficient for binding. We thus broadened our human proteome screening approach to accommodate these variations in the DDVF motif and screened the human proteome for the presence of either the D/E-x-V-F or the D/E-V-F motif. This screening identified 1526 human proteins (**Table S1B)** including 48 previously identified RSK interactors. Among these proteins, all four isoforms of Fibroblast Growth Factor Receptor (FGFR1-4) beared an evolutionarily well-conserved DSVF motif in their C-terminal tails (**Figure 2A**) even though this region exhibits poor conservation with only 16 out of 61 conserved residues. Given that a previous study reported an interaction between RSK2 and the C-terminal tail of the FGFR1 receptor (4), we sought to investigate whether this DSVF motif could mediate the interaction of FGFR1 with the KAKLGM site of RSK. The DSVF motif of all FGFR isoforms successfully docked into the RSK KAKLGM site in 5 out of 5 AlphaFold-multimer predictions (see **Supporting Figure S1C and Table S1D**). To experimentally assess the importance of the DSVF motif for the RSK:FGFR1 interaction, we conducted co-immunoprecipitation assays in HeLa cells co-transfected with plasmids expressing HA-FGFR1 (DSVF or DSVA mutant) and FLAG-RSK1 (wild type KAKLGM or KSEPPY mutant) (**Figure 2B**). Our data show that the DSVF motif of FGFR1 indeed interacts with the KAKLGM site of RSK1.

**Figure 2:**
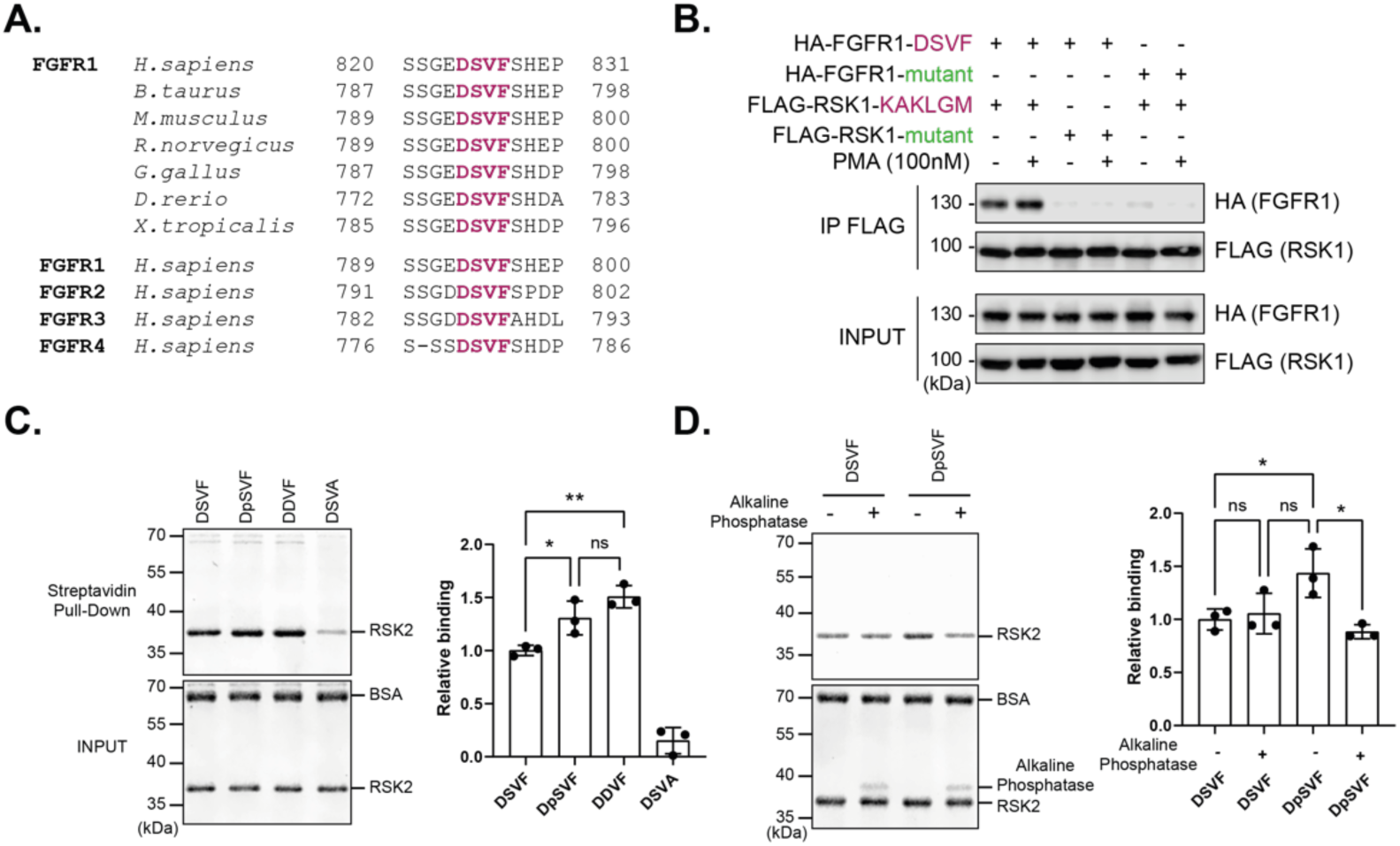
FGFR1 interacts with RSK1 via a DSVF motif whose phosphorylation increases interaction. **A.** The DSVF motif from FGFR1 is highly conserved in all 4 FGFR isoforms and across evolution. **B.** Co-immunoprecipitation of HA-FGFR1 DSVF or the DSVA mutant with FLAG-RSK1 KAKLGM or the KSEPPY mutant, from HeLa cells co-transfected with plasmids expressing indicated proteins. (INPUT = cell lysates; IP = immunoprecipitation), (n = 3). **C.** Assessment of the interaction between biotinylated peptides derived from FGFR1, bearing DSVF, DDVF, DSVA, or DpSVF (phosphoserine) motifs, and purified RSK2 (residues 44-367) using streptavidin pull-down assays (n = 3). **D.** Same experiment as in C comparing the pull-down efficiency of DSVF and DpSVF peptides on RSK2 with or without treatment of the peptides with alkaline phosphatase (n = 3). **C-D.** Quantitative analyses show the mean +/− SD of relative RSK pull-down efficiencies calculated from three independent experiments; P-values: *, p≤ 0.05; **, p≤0.01; ***, p≤0.001.

Interestingly, in the DSVF SLiM found in FGFR1, a serine residue is substituted for glutamate or aspartate negatively charged residues found in pathogens’ SLiMs. Since serine phosphorylation would restore a negative charge, we tested if the phosphorylation of the serine in the D<u>S</u>VF motif could increase the DSVF:RSK interaction. In support of this idea, Nadratowska-Wesolowska *et al.* reported a reduced FGFR1:RSK2 interaction after mutation of this serine into an alanine (4).

We thus performed streptavidin pull-down assays using biotinylated synthetic peptides derived from the FGFR1 sequence, with variation in the DSVF motif (DSVF, DpSVF (phosphoserine), DDVF, or DSVA) in the presence of a bacterially expressed, recombinant RSK2 (**Figure 2C**). As in the case of the full-length FGFR1 protein (**Figure 2B**), the DSVF-bearing peptide interacted better with RSK2 compared to the DSVA control (**Figure 2C**). The DDVF-bearing peptide exhibited even stronger binding to RSK2 than its wild type DSVF counterpart, suggesting that the introduction of a negative charge increased interaction. In line with this, we observed that the phosphopeptide DpSVF displayed a slightly but significantly stronger binding to RSK than the non-phosphorylated DSVF peptide (**Figure 2C**). Of note, the DpSVF peptide exhibited slightly lower RSK binding ability than the DDVF peptide. This lower binding of the DpSVF peptide to RSK2 might be attributed to the lower purity of the phosphorylated peptide yielded during synthesis when compared to the DDVF peptide, as assessed by routine mass spectrometry.

To confirm the impact of serine phosphorylation on binding, another set of pull-down assays was performed, comparing pull-down efficacies with and without alkaline phosphatase treatment of the peptides (**Figure 2D**). Again, a slight but significant difference was observed, showing that de-phosphorylation of the serine residue in the DpSVF phosphopeptide decreased RSK binding. These data suggest the existence of an additional layer of regulation for RSK binding, where kinases could modulate the binding affinity between FGFRs and RSKs through phosphorylation.

### 4. CNKSR2 also interacts with RSK through a DSVF motif

As the three RSK-interacting proteins identified above were known as effectors/regulators of the RAS-ERK MAPK pathway, we wondered whether additional protein from this pathway would use a similar SLiM to interact with RSK. From our 1526 human protein list (**Table S1B**) we selected Connector enhancer of kinase suppressor of ras 2 (CNKSR2) which harbors a well conserved DSVF motif (**Figure 3A**) predicted to dock in RSK KAKLGM D-site in 5 out of 5 models (**Table S1D**). Again, co-immunoprecipitations assays performed in HEK293T cells co-transfected with plasmids expressing FLAG-CNKSR2 (DSVF or DSVA mutant) and HA-RSK1 (KAKLGM or KSEPPY mutant) showed that the DSVF motif of CNKSR2 interacts with the KAKLGM site of RSK1 (**Figure 3B**).

**Figure 3:**
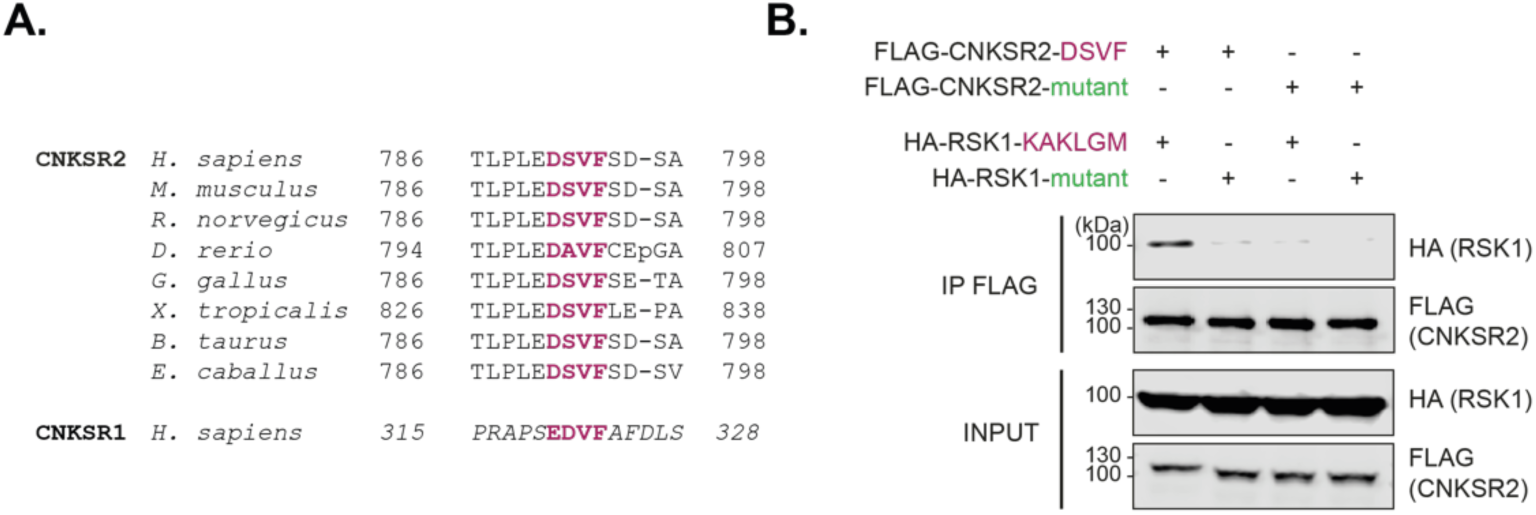
CNKSR2 interacts with RSK1 through a DSVF motif. **A.** The DSVF motif from CNKSR2 is conserved in CNKSR1 and across evolution. **B.** Co-immunoprecipitation of FLAG-CNKSR2 DSVF or the DSVA mutant with FLAG-RSK1 KAKLGM or the KSEPPY mutant, from HEK293T cells co-transfected with plasmids expressing indicated proteins. (n = 2).

### 5. The RSK SLiM-binding site participates in the regulation of the MAPK pathway and RSK itself

Cellular RSK interactors identified above, such as SPRED2 (28), GAB3 (14),FGFR1 (4), CNKSR2 (29), are related to the RAS-ERK MAP kinase pathway, as does SOS1, which was suggested to interact with the same RSK SLiM-binding site (5, 30). Interaction between these proteins and RSKs might thus be involved in fine tuning the RAS-ERK-MAPK pathway (**Figure 4A-B**). To test this hypothesis, we monitored ERK activation upon bFGF stimulation in cells expressing either RSK wild-type (WT) or the KAKLGM-to-KSEPPY mutant. For this investigation, we made use of our previously generated HeLa M RSK1/2 double KO cells (RSK-DKO), which minimally express, RSK3 and RSK4 mRNAs, and are thus virtually RSK-KO cells (13). These cells were transduced with lentiviral vectors expressing either RSK1 WT (KAKLGM) or the KSEPPY mutant (**Figure 4C**). Upon bFGF stimulation, the phospho-ERK signal remained more elevated in RSK-DKO cells transduced with the empty vector than in cells transduced with the RSK1 WT (KAKLGM) expression vector (**Figure 4D-E**), thus corroborating a role for RSK in the negative feedback of the RAS-MAPK pathway. Furthermore, cells expressing the RSK1 KSEPPY mutant exhibited increased phospho-ERK levels over time compared to cells expressing WT RSK1 (**Figure 4D-E**). This observation suggests that the DDVF-like SLiM-mediated interaction of cellular proteins with RSKs contributes to the overall negative feedback response of the RAS-ERK MAP kinase pathway.

**Figure 4:**
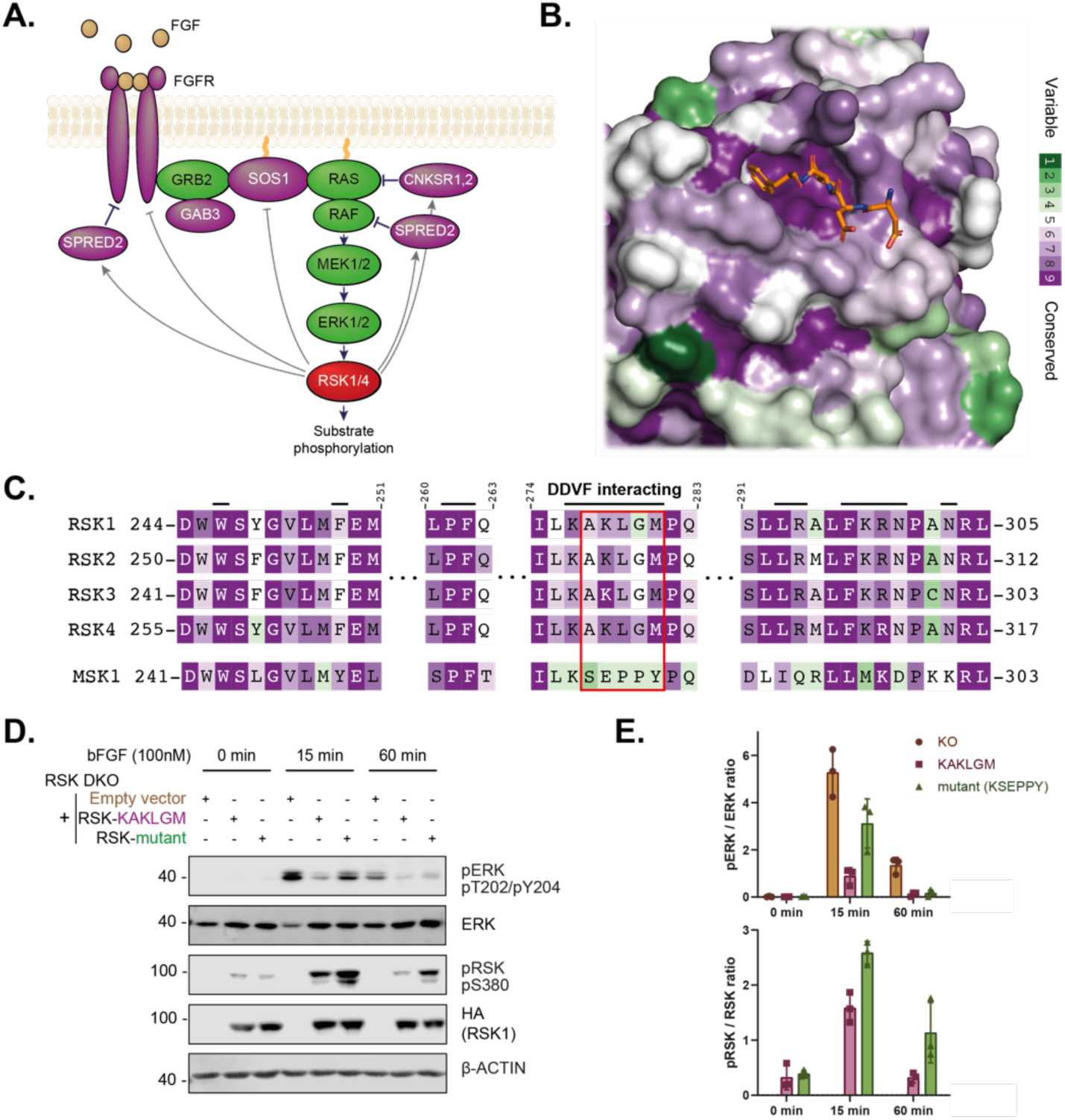
The conserved DDVF-interacting region contributes to an RSK-mediated negative feedback of the ERK-MAPK pathway in HeLa cells. **A.** Several proteins from the RAS-MAPK pathway bear a DDVF-like motif (purple) (31). In the case of SPRED2, FGFR1 and SOS1, indicated negative feedbacks were described in (4–6, 28). **B-C.** RSK tridimensional structure (B) and sequence alignment (C) showing the conservation of the RSK DDVF docking site across evolution (from purple = highly conserved to green = poorly conserved). **B.** RSK2 tridimensional structure colored according to conservation across evolution with the DDVF peptide colored in orange (PDB: 7OPO, (14)). **C.** Sequence alignment showing that the DDVF-interacting residues (indicated by upper black bars) in the KAKLGM region are well conserved across all four RSK isoforms but not in the closely related MSK1, justifying the KAKLGM-to-KSEPPY mutation. **D-E.** Immunoblotting of lysates from RSK-DKO cells transduced with an empty vector (orange), or with vectors expressing RSK1 KAKLGM WT (purple) or the KSEPPY mutant (green). Cells were starved for 14-16 hours and then stimulated with 100nM bFGF for indicated periods of time. Western blots (n = 3) were quantified to calculate phosphor-ERK/ERK and phosphor-RSK/RSK ratios (mean +/− SD).

## Discussion

SLiMs are emerging as omnipresent motifs which fine tune cellular signaling pathways through subtle protein-protein interactions. Given their small size (usually 3 to 12 amino acids) and unstructured nature, SliMs can easily arise from convergent evolution, making them appealing targets for pathogenic mimicry, in particular in the case of RNA viruses (22, 23). Previous studies (13, 14, 24) have revealed that unrelated pathogens, including RNA viruses, DNA viruses and bacteria expressed proteins carrying a DDVF-like SLiM enabling them to interact with the highly conserved KAKLGM SLiM-binding site of RSKs. We postulated that these pathogens creatively evolved to imitate cellular proteins, which bear a similar DDVF motif to influence RSK activity or subcellular localization through this docking site. To find such regulator proteins, we screened the human proteome for proteins containing an evolutionary conserved DDVF-like motif. AlphaFold-multimer provided an effective support to the candidates’ selection pipeline prior to in vivo validation, as the predictions displayed the expected interface in 5 out of 5 models for the 4 confirmed RSK-interacting proteins identified. In all cases, the DDVF-like SLiM of human proteins interacted with the KAKLGM region of RSK in the same way as that proposed for the SLiM of pathogens’ proteins. The similarity of the interfaces was structurally confirmed for RSK interaction with Kaposi sarcoma-associated herpes virus protein ORF45 (14) and with the human protein SPRED2 (6).

Our work shows that the DDVF-like SLiM occurring in pathogens’ proteins mimics a SLiM likely to be important for fine tuning of the RAS-ERK MAP kinase pathway. This pathway and the downstream RSK kinases phosphorylate a surprisingly large variety of substrates (32) and thereby regulate important biological processes including cell survival, growth and translation. Tight regulation of this pathway is crucial to prevent the disastrous consequences of its overactivation. Previous research has highlighted the role of RSK, SPRED2, as well as RSK targets such as FGFR1 and SOS1 in negative feedback to the ERK-MAPK pathway (7). We identified GAB3 and CNKSR2 as new RSK interactors. GAB proteins are recognized for their role in modulating various signaling pathways, including MAP kinase pathways (33). GAB3, the least studied among the GAB isoforms, is the only one possessing a DDVF-like motif, suggesting that it acts through a mechanism involving RSK (**Supporting Figure S1B**). CNKSR proteins are also known modulators of the RAS-ERK MAP kinase pathway (34). Both CNKSR isoforms, CNKSR1 and 2, contain an EDVF motif that docks with the RSK KAKLGM region, as predicted by AlphaFold.

The RSK interaction motifs detected in FGFR1 and CNKSR2 (DSVF) contain a serine instead of a negatively charged amino acid (Asp or Glu) found in the SLiM of pathogens’ proteins. Interestingly, according to its phosphorylation status, the DSVF motif in FGFR1 exhibits a tunable affinity for the electropositive KAKLGM motif of RSK, (**Figure 2 C-D**). This observation underscores a novel layer of RSK regulation, potentially involving other Ser/Thr kinases, which would modulate the temporality or the subcellular localization of RSK association with those proteins, via phosphorylation of the SLiM.

As suggested by our data (**Figure 4**), the occurrence of DDVF-like SLiMs in these proteins could act to downregulate the RAS-ERK pathway at multiple levels, safeguarding against its overactivation in cells. The importance of the KAKLGM SLiM binding site in the regulation of RSKs’ physiological activity likely offered an easy target to pathogens for SLiM mimicry. On the one hand, by expressing proteins containing a DDVF-like SLiM, pathogens likely compete with endogenous DDVF-bearing proteins for RSK binding, thereby affecting the physiological regulation of the RAS-ERK-RSK pathway. Such dysregulation may lead to effects such as increased translation or dysregulated apoptosis, which may benefit the pathogen. Competition by pathogens’ proteins is likely an effective mechanism in the case of viral infections given the often-high expression level of viral proteins in the infected cell.

On the other hand, as shown in for *Yersinia* and for cardioviruses, interaction of pathogens’ proteins with RSKs through SLiM mimicry may be used to retarget RSKs toward unconventional substrates. *Yersinia* YopM was shown to form a complex with several proteins including RSK, another kinase named PRK or PKN, as well as with pyrin, leading to pyrin phosphorylation and ultimately to inflammasome inhibition (35, 36). In the case of cardioviruses, L protein interaction was shown to target RSK toward the nuclear pore complex where RSK phosphorylates FG-nucleoporins such as NUP98, thereby disrupting nucleo-cytoplasmic trafficking in the cell (37). Pyrin and NUP98 were not detected among the regular RSK substrates (32), suggesting that YopM and L can act as bridging platforms to increase the contact between RSKs and unconventional substrates, thereby promoting their phosphorylation by RSKs.

The interface between DDVF-like SLiMs and RSKs offers a screening opportunity for molecules that modulate RSK functions or prevent their hijacking by pathogens, as shown in a recent screening setup (30) and in a study targeting Kaposi’s sarcoma-associated herpesvirus (KSHV) infection (38). However, it is worth noting that inhibition of the RSK-SLiM interaction in this context may in part compromise the fine regulation of the RAS-MAPK pathway. This raises concerns in the case of long-term treatments as the RAS-MAPK pathway is often hyperactivated in cancer. In contrast, molecules inhibiting DDVF-RSK interaction might prove useful as short term treatment to attenuate the severity of acute viral or bacterial infections.

## Material and Methods

### Cells

HEK293T (39) and HeLa M cells, a subclone of HeLa cells kindly provided by R. H. Silverman (40) referred to as HeLa cells, were maintained in Dulbecco’s Modified Eagle Medium (Lonza) supplemented with 10% fetal bovine serum (Sigma), 100 U/mL penicillin and 100μg/mL streptomycin (Lonza). RSK1 and RSK2 double knock out HeLa M cells referred to as RSK-DKO cells were generated and validated in (13). One transcribed RSK1 allele is however detected carrying a 81nt in-frame deletion. These cells express low levels, if any, of RSK3 and RSK4 mRNA and can thus be considered as virtually RSK-KO cells. All cells were cultured at 37°C in a humidified atmosphere containing 5% CO2.

### Plasmids, retroviral and lentiviral constructs

Expression plasmids and retro/lentiviral expression vectors are presented in **Table S2**. Note that tagged proteins expressed by these vectors contain either 3xFLAG or 3xHA tag which are referred to as FLAG- and HA-in the text and figures. Plasmid vectors include pcDNA3 (Invitrogen) for mammalian expression. Retroviral expression vectors were derived from pQCXIH (Clontech) and lentiviral vectors were derived from pTM952, a derivative of pCCLsin.PPT.hPGK.GFP.pre (41, 42). Lentiviral constructs expressing human RSKs were constructed using the Gateway technology (Invitrogen) from donor plasmids. pTM1116 encodes Human RSK1 (37). Donor plasmids encoding Hs.SPRED2 Hs.SPRED1 and HsCNKSR2 were kindly provided by Dominic Esposito through the Addgene collection (Addgene refs: #70573, #70607, #70605 and #70313 respectively), HsGAB3 through the DNASU collection (ref: HsCD00081299) and Hs.FGFR1 was kindly provided by Jean-Baptiste Demoulin. Note that, where indicated, lentiviral or retroviral constructs were transfected as expression plasmids instead of being transduced.

### Cell transfection and stimulation

HEK293T or HeLa cells, seeded the day before, were transfected using Lipofectamine 2000 (Thermofisher) according to manufacturer’s instructions with a ratio of 0.5 μg / 2 μl or 2.5 μg / 7.5 μl of DNA/transfection reagent for 24-well plates or 6-well plates, respectively. When indicated, cells were stimulated with 100nM Phorbol-12-myristate-13-acetate (PMA) (Sigma-Aldrich, #P8139) at 37°C for 20 minutes. In the case of bFGF treatments, cells were starved 6-8 hours after transfection for a period of 14-16 hours before stimulation with 100nM bFGF (Peprotech # 100-18B) and 3.33nM heparin at indicated timepoints.

### Immunoprecipitations

Transfected cells were lysed in lysis buffer (Tris-HCl 50 mM pH 8, NaCl 150 mM, NP40 0.5%, EDTA 2 mM, PMSF 1 mM and supplemented with protease/phosphatase inhibitors (Pierce)) and centrifugated at 14,000 x g for 10 min at 4°C. Cleared supernatants were incubated with anti-FLAG M2 Magnetic Beads (#M8823, Sigma-Aldrich) with gentle agitation for 4 hours at 4°C. Magnetic beads were then washed 3 times with the lysis buffer without inhibitors. Immunoprecipitated proteins were detected by Western-blot analysis.

### Western blot analysis

Total protein extracts were denatured for 5 min in Laemmli buffer at 95 °C, resolved by SDS-PAGE and transferred to polyvinylidene difluoride or nitrocellulose membranes (Immobilon; Millipore). Blocking of unspecific antigens was carried out in 5% nonfat dried milk in Tris-buffered saline (50 mM NaCl, 50 mM Tris-HCl, pH 7.5) for 1 hour at room temperature. Primary antibodies are listed in **Table S3.** Reference related to ERK antibody production can be found in (43). Primary antibodies were diluted in TBST (TBS containing 0.2% Tween 20) and incubated overnight at 4°C with gentle rolling. Membranes were washed three times in TBST for 5 min at room temperature. Species-matched IRDye or HRP secondary antibodies were diluted in blocking buffer as before and incubated at room temperature for 1h. Membranes were washed again three times in TBST for 5 min at room temperature. Fluorescent signal was detected through an Odyssey Fc infrared imaging system (Li-Cor) while HRP-generated signal was detected using Pierce SuperSignal or Cyanagen Supernova substrates on the same apparatus.

### RSK2 bacterial protein expression and purification

*E. coli* BL21-AI were transformed with pFB42, a plasmid encoding a 6-His N-Terminal fusion with the residues 44-367 of the murine RSK2 under the control of a T7 promoter upstream of the lac operator and constitutively expressing the LacI repressor. A single colony grown on an agar plate containing 250 ug/mL ampicillin was amplified in TSB medium supplemented with the same concentration of ampicillin and further grown under constant shaking at 37°C to an OD of about 0.6. IPTG and arabinose were then added at a final concentration of 1 mM and 0.2%, respectively, and cultures were incubated for another four hours at 37°C under agitation. Bacteria were harvested by centrifugation, resuspended in Tris-HCl 50 mM pH8, NaCl 150 mm, imidazole 20mM, 10 mg/mL lysozyme and supplemented with protease inhibitors (Roche) and further lysed by sonication. Cleared bacterial lysates were loaded on column packed with 1mL Ni-NTA agarose (QIAGEN) and washed with 5 column volumes (CV) of Tris-HCl 50 mM pH 8, NaCl 150 mM, imidazole 20 mM, followed by 5 CV of Tris-HCl 50 mM pH 8, NaCl 500 mM, imidazole 20 mM and eluted in 5 CV of Tris-HCl 50 mM pH 8, NaCl 150 mM, imidazole 300 mM. Eluted proteins were dialyzed overnight in Tris-HCl 50 mM pH 8, NaCl 150 mM supplemented with 10% glycerol. Protein purity (>90%) was assessed by SDS-PAGE analysis and Coomassie staining and protein concentration was measured by UV spectrophotometry with the extinction coefficient determined by ExPASy ProtParam (44). Aliquots of dialyzed RSK2 were snap-frozen and stored at −70°C until further use.

### Biotinylated FGFR1-derived peptide pull-down assay

For RSK2 pull-down assays, 400 pmol of biotinylated FGFR1-derived peptides were incubated in the presence or absence of 5 units of alkaline phosphatase (FastAP, #EF651, ThermoFischer Scientific) at 37°C in 10 mM Tris-HCl, 5 mM MgCl_2_, 100 mM KCl, 0.02% Triton X-100 and 100 ug/mL BSA in a total volume of 100 uL. Sequences of the biotinylated FGFR1-derived peptides were KDTRSSTCSSGE[DSVF]SHE with the DSVF region replaced by DSVA, DpSVF, DDVF or DSVA amino-acids as indicated. Mass spectrometry analysis confirmed the identity of the synthesized peptides. After one hour, 400 pmol of recombinant RSK2 (44–367) diluted in 100 uL of the same buffer supplemented with 2 mM imidazole and 1 mM Na3VO4 was mixed with FGFR1-derived biotinylated peptides and incubated for 2 hours at 4 °C under gentle end-to-end agitation. Final concentrations of both RSK2 and the FGFR1-derived peptides were 2 uM. After two hours, 20 uL of a 50% slurry of streptavidin magnetic beads (#88817, Pierce) were added and incubated as before. Bound peptides were then washed with three times with 50 mM Tris-HCl pH 8, 150 mM NaCl, 0.5% NP40, 2 mM EDTA and precipitated RSK2 was eluted in Laemmli buffer and analyzed by SDS-PAGE and Coomassie staining. Gel imaging and protein quantification were performed using an Odyssey Fc imaging system interfaced with Image Studio 5.2 (LI-COR).

### Alignments

Clustal Omega software (https://www.ebi.ac.uk/Tools/msa/clustalo/) was used to produce multiple sequence alignments from human SPRED1-3 (Uniprot sequences Q7Z699, Q7Z698, and Q2MJR0), FGFR1-4 (Uniprot sequences P11362, P21802, P22607, and P22455), and RSK1-4 (Uniprot sequences Q15418, P51812, Q15349, and Q9UK32). To calculate the percentage of conservation of the unstructured C-term of the FGFR1-4 proteins, region annotated as unstructured according to MobiDB (45) (https://mobidb.org/) were aligned with Clustal Omega. Multiple sequence alignments of SPRED2 and FGFR1 from various species were retrieved from the now discontinued Homologene database. Multiple sequence alignments of GAB3 and CNKSR2 from various species was provided by Gene - NCBI (https://www.ncbi.nlm.nih.gov/datasets/gene/). Multiple sequence alignments of RSK1-4 across evolution were given by Proviz (46) (http://proviz.ucd.ie/), and coloring scheme according to conservation was calculated by ConSurf web server (https://consurf.tau.ac.il/).

### AlphaFold-Multimer structure predictions and visualization

Protein complex structure predictions were generated on a locally-installed AlphaFold v2.3.2 (19) kindly implemented by Raphael Helaers from the de Duve Institute. Protein complex predictions were performed in the multimer mode using defaults parameters. Inputs were sequences derived from a fragment (amino acid 161-337) of the N-terminal kinase domain from the human RSK2 protein (accession number: P51812) and peptide sequences of 103 amino acid centered on the valine of the D/E-D/E-V-F motif except in situations where the motif was located at the N- or C-terminus of the protein candidates. All structural predictions and associated accuracy metrics have been made available in the zenodo open research data repository (Parts 1 and 2: doi:10.5281/zenodo.10630297, Parts 3 and 4: doi:10.5281/zenodo.10653847 and part 5: doi:10.5281/zenodo.10658285). Multiple structure alignment and visualization were generated using PyMOL 2.5.4 (Schrodinger, LLC).

### Quantification and statistical analysis

Statistical significance was determined using one-way *ANOVA* as implemented in the Prism 8.0.02 statistical analysis software (GraphPad Software, Inc., San Diego, CA). The number of independent experiments (n) and statistical comparison groups are indicated in the figures or figure legends. Asterisks denote the statistical significance of the indicated comparisons as follows: *, p≤ 0.05; **, p≤0.01; ***, p≤0.001.

## Acknowledgments

We thank Stéphane Messe for expert technical assistance, Anca Marian for her work as a student, Fabian Borghese for the construction of plasmid pFB. Finally, we thank Raphaël Helaers for the implementation of the AlphaFold-multimer structure prediction software on the high-performance computing infrastructure of the de Duve Institute.

## Supporting information captions

Table S1. **SLiMSearch and AlphaFold Screens**. **S1A**: Results of the D/E-D/E-V-F screening using SLiMsearch4, crossed with interaction data avaible on BioGrid, Intact, and PhosphositePlus. **S1B**: Results of the D/E-x-V-F and D/E-V-F screening crossed with interaction data mentioned hereabove. **S1C**: Prediction of motif docking within the KAKLGM pocket of RSK using AlphaFold multimer. Motifs are derived from proteins in the S1A section. **S1D**: Prediction of FGFRs/CNKSR2 DSVF motifs docking within the KAKLGM pocket of RSK using AlphaFold multimer.

Table S2. **Plasmids used.**

Table S3. **Antibodies used.**

Figure S1. **Comparison of SPRED/GAB isoforms sequences and AlphaFold docking.**

**A-B**. Conservation of SPRED2 DDVF (purple) motif across all three SPRED (**A**) and GAB (**B**) proteins. **C.** AlphaFold multimer docks DDVF-like motif from SPRED2 (green), GAB3 (red), FGFR1-4 (blue), and CNKSR2 (purple) in RSK KAKLGM docking site. The obtained structures show the same pose found in the crustal structure of ORF45 DDVF (orange) bound to RSK2 (PDB 7opo).

## References

1. Kumar R, Khandelwal N, Thachamvally R, Tripathi BN, Barua S, Kashyap SK, et al. Role of MAPK/MNK1 signaling in virus replication. Virus Res. 2018;253:48–61.

2. Pleschka S. RNA viruses and the mitogenic Raf/MEK/ERK signal transduction cascade. Biol Chem. 2008;389(10):1273–82.

3. Romeo Y, Zhang X, Roux PP. Regulation and function of the RSK family of protein kinases. Biochem J. 2012;441(2):553–69.

4. Nadratowska-Wesolowska B, Haugsten EM, Zakrzewska M, Jakimowicz P, Zhen Y, Pajdzik D, et al. RSK2 regulates endocytosis of FGF receptor 1 by phosphorylation on serine 789. Oncogene. 2014;33(40):4823–36.

5. Saha M, Carriere A, Cheerathodi M, Zhang X, Lavoie G, Rush J, et al. RSK phosphorylates SOS1 creating 14-3-3-docking sites and negatively regulating MAPK activation. Biochem J. 2012;447(1):159–66.

6. Lopez J, Bonsor DA, Sale MJ, Urisman A, Mehalko JL, Cabanski-Dunning M, et al. The ribosomal S6 kinase 2 (RSK2)-SPRED2 complex regulates the phosphorylation of RSK substrates and MAPK signaling. J Biol Chem. 2023;299(6):104789.

7. Fricke AL, Mühlhäuser WWD, Reimann L, Zimmermann JP, Reichenbach C, Knapp B, et al. Phosphoproteomics Profiling Defines a Target Landscape of the Basophilic Protein Kinases AKT, S6K, and RSK in Skeletal Myotubes. J Proteome Res. 2023;22(3):768–89.

8. Hebron KE, Hernandez ER, Yohe ME. The RASopathies: from pathogenetics to therapeutics. Dis Model Mech. 2022;15(2).

9. Arthur JS, Ley SC. Mitogen-activated protein kinases in innate immunity. Nat Rev Immunol. 2013;13(9):679–92.

10. Hoshino R, Chatani Y, Yamori T, Tsuruo T, Oka H, Yoshida O, et al. Constitutive activation of the 41-/43-kDa mitogen-activated protein kinase signaling pathway in human tumors. Oncogene. 1999;18(3):813–22.

11. Romeo Y, Roux PP. Paving the way for targeting RSK in cancer. Expert Opin Ther Targets. 2011;15(1):5–9.

12. Youn M, Gomez JO, Mark K, Sakamoto KM. RSK Isoforms in Acute Myeloid Leukemia. Biomedicines. 2021;9(7).

13. Sorgeloos F, Peeters M, Hayashi Y, Borghese F, Capelli N, Drappier M, et al. A case of convergent evolution: Several viral and bacterial pathogens hijack RSK kinases through a common linear motif. Proc Natl Acad Sci U S A. 2022;119(5).

14. Alexa A, Sok P, Gross F, Albert K, Kobori E, Poti AL, et al. A non-catalytic herpesviral protein reconfigures ERK-RSK signaling by targeting kinase docking systems in the host. Nat Commun. 2022;13(1):472.

15. Delaunoy JP, Dubos A, Marques Pereira P, Hanauer A. Identification of novel mutations in the RSK2 gene (RPS6KA3) in patients with Coffin-Lowry syndrome. Clin Genet. 2006;70(2):161–6.

16. Reményi A, Good MC, Lim WA. Docking interactions in protein kinase and phosphatase networks. Curr Opin Struct Biol. 2006;16(6):676–85.

17. Benz C, Ali M, Krystkowiak I, Simonetti L, Sayadi A, Mihalic F, et al. Proteome-scale mapping of binding sites in the unstructured regions of the human proteome. Mol Syst Biol. 2022;18(1):e10584.

18. Rrustemi T, Meyer K, Roske Y, Uyar B, Akalin A, Imami K, et al. Pathogenic mutations of human phosphorylation sites affect protein-protein interactions. bioRxiv. 2023:2023.08.01.551433.

19. Evans R, O’Neill M, Pritzel A, Antropova N, Senior A, Green T, et al. Protein complex prediction with AlphaFold-Multimer. bioRxiv. 2022:2021.10.04.463034.

20. Verburgt J, Zhang Z, Kihara D. Multi-level analysis of intrinsically disordered protein docking methods. Methods. 2022;204:55–63.

21. Lee CY, Hubrich D, Varga JK, Schäfer C, Welzel M, Schumbera E, et al. Systematic discovery of protein interaction interfaces using AlphaFold and experimental validation. Mol Syst Biol. 2024;20(2):75–97.

22. Mihalič F, Simonetti L, Giudice G, Sander MR, Lindqvist R, Peters MBA, et al. Large-scale phage-based screening reveals extensive pan-viral mimicry of host short linear motifs. Nat Commun. 2023;14(1):2409.

23. Glavina J, Palopoli N, Chemes LB. Evolution of SLiM-mediated hijack functions in intrinsically disordered viral proteins. Essays Biochem. 2022;66(7):945–58.

24. Hentschke M, Berneking L, Belmar Campos C, Buck F, Ruckdeschel K, Aepfelbacher M. Yersinia virulence factor YopM induces sustained RSK activation by interfering with dephosphorylation. PLoS One. 2010;5(10).

25. Krystkowiak I, Davey NE. SLiMSearch: a framework for proteome-wide discovery and annotation of functional modules in intrinsically disordered regions. Nucleic Acids Res. 2017;45(W1):W464–w9.

26. Del Toro N, Shrivastava A, Ragueneau E, Meldal B, Combe C, Barrera E, et al. The IntAct database: efficient access to fine-grained molecular interaction data. Nucleic Acids Res. 2022;50(D1):D648–d53.

27. Oughtred R, Rust J, Chang C, Breitkreutz BJ, Stark C, Willems A, et al. The BioGRID database: A comprehensive biomedical resource of curated protein, genetic, and chemical interactions. Protein Sci. 2021;30(1):187–200.

28. Mardakheh FK, Yekezare M, Machesky LM, Heath JK. Spred2 interaction with the late endosomal protein NBR1 down-regulates fibroblast growth factor receptor signaling. J Cell Biol. 2009;187(2):265–77.

29. Ito H, Nagata KI. Functions of CNKSR2 and Its Association with Neurodevelopmental Disorders. Cells. 2022;11(2).

30. Póti Á L, Dénes L, Papp K, Bató C, Bánóczi Z, Reményi A, et al. Phosphorylation-Assisted Luciferase Complementation Assay Designed to Monitor Kinase Activity and Kinase-Domain-Mediated Protein-Protein Binding. Int J Mol Sci. 2023;24(19).

31. Anjum R, Blenis J. The RSK family of kinases: emerging roles in cellular signalling. Nat Rev Mol Cell Biol. 2008;9(10):747–58.

32. Lizcano-Perret B, Vertommen D, Herinckx G, Calabrese V, Gatto L, Roux PP, et al. Identification of RSK substrates using an analog-sensitive kinase approach. J Biol Chem. 2024;300(3):105739.

33. Liu Y, Rohrschneider LR. The gift of Gab. FEBS Lett. 2002;515(1-3):1–7.

34. Udaykumar N, Zaidi MAA, Rai A, Sen J. CNKSR2, a downstream mediator of retinoic acid signaling, modulates the Ras/Raf/MEK pathway to regulate patterning and invagination of the chick forebrain roof plate. Development. 2023;150(3).

35. McDonald C, Vacratsis PO, Bliska JB, Dixon JE. The yersinia virulence factor YopM forms a novel protein complex with two cellular kinases. J Biol Chem. 2003;278(20):18514–23.

36. Park YH, Remmers EF, Lee W, Ombrello AK, Chung LK, Shilei Z, et al. Ancient familial Mediterranean fever mutations in human pyrin and resistance to Yersinia pestis. Nat Immunol. 2020;21(8):857–67.

37. Lizcano-Perret B, Lardinois C, Wavreil F, Hauchamps P, Herinckx G, Sorgeloos F, et al. Cardiovirus leader proteins retarget RSK kinases toward alternative substrates to perturb nucleocytoplasmic traffic. PLoS Pathog. 2022;18(12):e1011042.

38. Li X, Huang L, Xiao Y, Yao X, Long X, Zhu F, et al. Development of an ORF45-Derived Peptide To Inhibit the Sustained RSK Activation and Lytic Replication of Kaposi’s Sarcoma-Associated Herpesvirus. J Virol. 2019;93(10).

39. DuBridge RB, Tang P, Hsia HC, Leong PM, Miller JH, Calos MP. Analysis of mutation in human cells by using an Epstein-Barr virus shuttle system. Mol Cell Biol. 1987;7(1):379–87.

40. Dong B, Niwa M, Walter P, Silverman RH. Basis for regulated RNA cleavage by functional analysis of RNase L and Ire1p. Rna. 2001;7(3):361–73.

41. Cesaro T, Hayashi Y, Borghese F, Vertommen D, Wavreil F, Michiels T. PKR activity modulation by phosphomimetic mutations of serine residues located three aminoacids upstream of double-stranded RNA binding motifs. Sci Rep. 2021;11(1):9188.

42. Follenzi A, Ailles LE, Bakovic S, Geuna M, Naldini L. Gene transfer by lentiviral vectors is limited by nuclear translocation and rescued by HIV-1 pol sequences. Nat Genet. 2000;25(2):217–22.

43. Leevers SJ, Marshall CJ. Activation of extracellular signal-regulated kinase, ERK2, by p21ras oncoprotein. Embo j. 1992;11(2):569–74.

44. Wilkins MR, Gasteiger E, Bairoch A, Sanchez JC, Williams KL, Appel RD, et al. Protein identification and analysis tools in the ExPASy server. Methods Mol Biol. 1999;112:531–52.

45. Piovesan D, Del Conte A, Clementel D, Monzon AM, Bevilacqua M, Aspromonte MC, et al. MobiDB: 10 years of intrinsically disordered proteins. Nucleic Acids Res. 2023;51(D1):D438–d44.

46. Jehl P, Manguy J, Shields DC, Higgins DG, Davey NE. ProViz-a web-based visualization tool to investigate the functional and evolutionary features of protein sequences. Nucleic Acids Res. 2016;44(W1):W11–5.

