## Supplementary material for "The “DDVF” motif used by viral and bacterial proteins to hijack RSK kinases evolved as a mimic of a short linear motif (SLiM) found in proteins related to the RAS-ERK MAP kinase pathway": Figure S1

**A.**

```
hSPRED3   48  YVIHGERLRDQKTTLECTLKPGLVYNKVNPIFHHWSLGDCKFGLTFQSPAEADEFQKSLL 107
hSPRED1   57  FFIRGERLRDKMVVLECMLKKDLIYNKVTPTFHHWKIDDKKFGLTFQSPADARAFDRGIR 117
hSPRED2   56  FLIHGERQKDKLVVLECYVRKDLVYTKANPTFHHWKVDNRKFGLTFQSPADARAFDRGVR 116
              :.*:*** :*: ..*** :: .*:*.*..* ****.:.: **********:*  *::.:

hSPRED3   106 AALAALGRGSLTPSSSSSSSSPSQDTAETPCP--LTSHV-----DSDS--SSSHSRQETP 158
hSPRED1   116 RAIEDISQGCPESKNEAEGADDLQANEEDSSSSLVKDHLFQQETVVTSEPYRSSNIRPSP 177
hSPRED2   115 KAIEDLIEGSTTSSSTIHNEAELGD-----------DDVFTTATDSSS---NSSQKREQP 162
               *:  : .*.   ..   .                 ..:        *    * . :  *
```

**B.**

```
hGAB3      86  FIVKTTSRTFYLVAKTEQEMQVWVHSISQVCNLGHLEDGADSMESLSYTPSSLQPSSAS- 144
hGAB1      85  FDINTIDRIFYLVADSEEEMNKWVRCICDICGFNPTEEDPVKPPGS--SLQA----PADL 138
hGAB2      86  FDIKTSERTFYLVAETEEDMNKWVQSICQICGFNQAEESTDSLRNVSSAGHGPRSSPAEL 145
hGAB4      121 FDIKTSERTFYLVAETREDMNEWVQSICQICGFRQE-ESTGFLGNISSASHGLCSSPAEP 179
               * ::* .* *****.:.::*: **:.*.::*.:    :.     .   :  .     *.

hGAB3      145 -----SLLTAH-------AASSSLPRDDPNTNAVA-TEETRSESELLFLPDYLVLSNCET 191
hGAB1      192 PLA-------INTAPPSTQADSSSATLPPPYQLINVPPHLETLGIQEDPQDYLLLINCQS 191
hGAB2      201 SSSSQHLLRERKSSAPSHSSQPTLFTFEPPV-----SNHMQPTLSTSAPQEYLYLHQCIS 200
hGAB4      223 SCSHQHLPQEQEP------------TSEPPV-----SHCVPPTWPIPAPPGCLRSHQHAS 222
                                           *                       *   :  :

hGAB3      192 GRLHHTSLPTRCDSWSNSDRSLEQASFDDVFVDCLQPLPSSHLVH---------PSCHGS 242
hGAB1      192 KKPEPTR----THADSAKSTS--------SETDCNDNVPSHKNPASSQSKHGMNGFFQQQ 239
hGAB2      201 RRAENAR----SASFSQGTRA--------SFLMRSDTAVQKLAQGNGHCVNGISGQVHG- 247
hGAB4      223 QRAEHAR----SASFSQGSEA--------PFIMRRNTAMQNLAQHSGYSVDGVSGHIHG- 269
```

**C.**

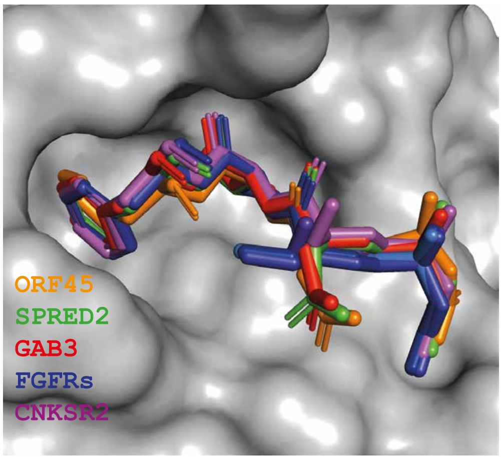
